# Myelin development in visual scene-network tracts beyond late childhood: A multimethod neuroimaging study

**DOI:** 10.1101/662809

**Authors:** Tobias W. Meissner, Erhan Genç, Burkhard Mädler, Sarah Weigelt

## Abstract

The visual scene-network—comprising the parahippocampal place area (PPA), retrosplenial cortex (RSC), and occipital place area (OPA)—shows a prolonged functional development. Structural development of white matter that underlies the scene-network has not been investigated despite its potential influence on scene-network function. The key factor for white matter maturation is myelination. However, research on myelination using the gold standard method of post-mortem histology is scarce. *In vivo* alternatives diffusion-weighed imaging (DWI) and myelin water imaging (MWI) so far report broad-scale findings that prohibit inferences concerning the scene-network. Here, we combine MWI, DWI tractography, and fMRI to investigate myelination in scene-network tracts in middle childhood, late childhood, and adulthood. We report increasing myelin from middle childhood to adulthood in right PPA-OPA, and trends towards increases in the left and right RSC-OPA tracts. Investigating tracts to regions highly connected with the scene-network, such as early visual cortex and the hippocampus did not yield any significant age group differences. Our findings indicate that structural development coincides with functional development in the scene-network, possibly enabling structure-function interactions.

## 1 Introduction

The human cortical visual system contains three high-level areas that preferentially respond to scenes compared to other stimuli, e.g. objects or faces: the parahippocampal place area (PPA, Epstein & Kanwisher, 1998), the retrosplenial cortex (RSC, O’Craven & Kanwisher, 2000), and the occipital place area (OPA, Grill-Spector, 2003; Hasson, Harel, Levy, & Malach, 2003). This functional network is strongly involved in scene processing (Bettencourt & Xu, 2013; Dilks, Julian, Kubilius, Spelke, & Kanwisher, 2011; Epstein, Higgins, Jablonski, & Feiler, 2007), but also in orientation and navigation (Epstein, 2008; Julian, Ryan, Hamilton, & Epstein, 2016). The scene-network’s components are already evident in middle childhood but at least for the PPA and the OPA there is evidence for a protracted development in terms of functional size and scene-selectivity beyond late childhood, possibly until adulthood (Chai, Ofen, Jacobs, & Gabrieli, 2010; Golarai et al., 2007; Meissner, Nordt, & Weigelt, 2019).

Despite this commencing understanding of the developmental trajectory of scene-network function between middle childhood and adulthood, the development of the white matter structure underlying the scene-network has not received attention so far. In other brain areas, white matter microstructure changes were shown to be an underlying mechanism for specific cognitive development or differences, as has been evidenced for musical proficiency (Bengtsson et al., 2005), vocabulary development (Pujol et al., 2006), and many other cognitive abilities (for an overview see Fields, 2008). Thus, maturational status of scene-network white matter structure might influence scene-network gray matter functional development, or vice versa (Fields, 2015; Zatorre, Fields, & Johansen-Berg, 2012).

The PPA, RSC, and OPA contribute to the complex tasks of scene processing and navigation through (at least partially) distinct functional response properties. In brief, the PPA is specialized on spatial layout detection and categorization of scenes; the RSC is specialized on cognitive mapping of the current view into an allocentric environment/reference frame and the encoding & retrieval of spatial & semantic scene context; and the OPA is specialized on identifying navigation possibilities in scenes as well as representing first-person perspective spatial relations within scenes (e.g. Baldassano, Esteva, Fei-Fei, & Beck, 2016; Epstein & Higgins, 2007; Epstein, Parker, & Feiler, 2007; Hutchison, Culham, Everling, Flanagan, & Gallivan, 2014; Vass & Epstein, 2013). Due to these distinct contributions, an efficient and mature transmission of signals between these areas is considered crucial for building an integrated perception of scenes. Further, the scene-network does not work in an isolated fashion. On the one hand, following the hierarchical organizational principle of the visual cortex, the early visual cortex (EVC) is a major input area to the PPA, RSC, and OPA (Grill-Spector & Malach, 2004). Unsurprisingly, studies in the past decades show that scene-selective areas are retinotopically organized and show strong functional connectivity to the EVC (Baldassano, Esteva et al., 2016; Baldassano, Fei-Fei, & Beck, 2016; Epstein & Baker, 2019). On the other hand, recent evidence suggests that the hippocampus (HC) might be part of the scene-network or at least a major input-output region (Baldassano, Beck, & Fei-Fei, 2013; Baldassano, Esteva et al., 2016; Dalton, Zeidman, McCormick, & Maguire, 2018; Graham, Barense, & Lee, 2010; Hodgetts et al., 2017; Hodgetts, Shine, Lawrence, Downing, & Graham, 2016; Zeidman, Mullally, & Maguire, 2015). Consequently, an efficient signal transmission between the scene-network areas and key areas working in concert with them to achieve scene perception should be an important developmental step.

Signal transmission can be optimized through increasing speed, synchrony, or reliability—all of which are mediated by increases in axon myelination (E. M. Miller, 1994; Zatorre et al., 2012). Axon myelination has traditionally been measured in post-mortem histological studies. However, post mortem-studies are rare in general and most studies focus on newborns’ and young infants’ gray matter myelin content. In the only (to the best of our knowledge) histological study investigating white matter myelin development beyond middle childhood, the authors report tract-specific maturation patterns featuring peak myelin growth rates within the first two years after birth as well as continued maturation up middle childhood (Yakovlev & Lecours, 1967). Evidence for development beyond childhood was limited to intracortical neuropil and association areas but should be regarded as rather anecdotal due to the low number of investigated tracts and specimens in that age group.

Due to the very limited availability of specimens for post-mortem histological myelin assessment, the advance of diffusion-weighed imaging (DWI), a non-invasive magnetic resonance imaging (MRI) method that has the potential to inform about myelin *in vivo*, represented a milestone. DWI has since been applied to probe developmental changes in white matter myelination extensively. However, most studies focused on major long fiber tracts, such as the internal capsule or the corticospinal tract, that can be readily identified (semi-) automatically using brain atlases (Lebel & Beaulieu, 2011; Mukherjee et al., 2001). As most long tracts are not directly involved in the visual scene-processing system and effects of age on white matter maturation were shown to be tract-specific (Rollins et al., 2010), the current literature is not informative on scene-network white matter development. Short-range tracts, which are crucial for relaying information in specialized functional networks over short distances, such as the scene-network, are understudied. The only relevant findings that we found suggest ongoing myelination in temporal and parietal lobe short-range tracts (Oyefiade et al., 2018) or in white matter adjacent to dorsal and ventral visual stream cortical areas (Barnea-Goraly et al., 2005; Loenneker et al., 2011) and thus remain too unspecific for any inference on scene-network developmental trajectories.

DWI’s sensitivity to myelin stems from its sensitivity to the diffusion of water: Because myelin reduces the inter-axonal space, it reduces the diffusion of water perpendicular to the axon orientation, i.e. increasing diffusion anisotropy (Feldman, Yeatman, Lee, Barde, & Gaman-Bean, 2010). However, several microstructural properties, such as axon diameter (Takahashi et al., 2002), axon packing density (Takahashi et al., 2002), axon membrane permeability (Ford, Hackney, Lavi, Phillips, & Patel, 1998), and fiber geometry (van Wedeen, Hagmann, Tseng, Reese, & Weisskoff, 2005) affect diffusion tensor imaging (DTI) parameters, too. Therefore, deducing myelination or maturational status from DTI parameters alone is challenging in most cases (Jones, Knösche, & Turner, 2013).

Myelin water imaging (MWI, MacKay et al., 1994), another MRI technique, is sensitive to myelin, highly reproducible (Meyers et al., 2009), and not affected by other microstructural changes, (Laule et al., 2006; Laule et al., 2008; Moore et al., 2000). Thus, it gives a more direct estimation of the status of myelination than interpretation of DTI parameters alone. Yet, MWI has only recently become available in pediatric research settings thanks to advances in sequence design that drastically sped up acquisition time (for multi-component driven equilibrium single pulse observation of T1 and T2 (mcDESPOT) MWI see Deoni, Rutt, Arun, Pierpaoli, & Jones, 2008, for 3D multi-echo (ME) gradient spin echo (GRASE) MWI see Prasloski et al., 2012).

A series of MWI studies using mcDESPOT investigated infants and young children and found steep increases of myelin from birth to age two and a moderate increase thereafter (Dean et al., 2014; Dean et al., 2015; Deoni et al., 2011; Deoni, Dean, O’Muircheartaigh, Dirks, & Jerskey, 2012; Deoni, Dean, Remer, Dirks, & O’Muircheartaigh, 2015). Recent findings based on mcDESPOT and 3D ME GRASE MWI indicate that while myelin does not seem to increase between middle and late childhood, a pronounced increase of myelin occurs in adolescence in major white matter tracts (Geeraert et al., 2018; Meissner, Genç, Mädler, & Weigelt, 2019). However, scene-network specific data has not been analyzed until now.

To complement recent findings regarding the functional development of scene-network regions PPA, RSC, and OPA (Meissner, Nordt, & Weigelt, 2019), we combined MWI and DWI-based probabilistic tractography to probe the structural maturation, i.e. myelin water fraction (MWF), of white matter that underlies scene-network function in middle childhood (7-8 years), late childhood (11-12 years), and adulthood (19-24 years). As previous behavioral studies identified a marked improvement in the performance in scene processing around the age of 10 (Day, 1975; Mackworth & Bruner, 1970; Munsinger & Gummerman, 1967; Vurpillot, 1968), these age groups were specifically chosen to capture the neural status—possibly underlying the behavior—before and after the change, as well as in a mature reference group. Further, we tested whether tracts that connect the scene-network with their key input/output areas such as the EVC or the HC, show increased myelination over time. In an extended analysis, we tested whether DWI parameters mirror our MWI results.

## 2 Methods

### 2.1 Participants

We analyzed data of 18 children aged 7-8 (*Mean (M)* = 7.56, *standard deviation (SD)* = 0.51; 7 female; henceforth: 7-8yo), 13 children aged 11-12 (*M* = 11.23, *SD* = 0.44; 8 female; henceforth: 11-12yo) and 16 adults aged 19-24 (*M* = 20.69, *SD* = 1.14; 7 female) for this study. The original sample included one additional 7-8yo that was excluded due to severely impaired data quality in the DWI scan, one 7-8yo that did not complete the myelin water imaging scan, one 11-12yo, in which our localizer failed to reveal any scene-selective ROIs, and one further child whose age was mixed up and thus did not fit within the age group of either 7-8-yo or 11-12-yo. Sample size was determined by following standard sample sizes in the field of cross-sectional developmental neuroimaging studies (e.g. Golarai et al., 2007). Our study worked towards answering several associated research questions and included multiple MRI sequences. Thus, most participants’ localizer data (see 2.2.3 fMRI data processing and functional ROI definition of PPA, RSC, and OPA) was analyzed in a previous publication, which also holds detailed information on recruitment and compensation (Meissner, Nordt, & Weigelt, 2019). None of the following exclusion criteria, which were established prior to any data analysis, were fulfilled by any participant: visual impairments that could not be corrected-to-normal during MRI scanning, permanent make-up, tattoos on the head or the neck, irremovable metallic body jewelry, or other metallic parts inside the body (e.g. braces, surgical steel), preterm birth, past or current neurological or psychiatric conditions, pre-clinical fear of narrow spaces, chronic neck- or back-pain, circulatory or respiratory conditions that would make MRI scanning potentially unsafe (e.g. the potential need for an asthma inhaler within less than 30 s), body weight exceeding 100 kg, structural brain abnormalities.

### 2.2 Neuroimaging

All magnetic resonance images were acquired at the Neuroimaging Centre of the Research Department of Neuroscience at Ruhr University Bochum’s teaching hospital Bergmannsheil on a 3.0T Achieva scanner (Philips, Amsterdam, The Netherlands) using a 32-channel head coil. Acquisition of data reported in this manuscript was part of a longer protocol that included further functional scans. To reassure children and parents as well as to provide the possibility for low-threshold contact, children were accompanied by one of the experimenters in the scanner room throughout the entire procedure. Children who had not participated in an MRI study before were accustomed to the scanning environment, experimental procedure, and localizer task in a custom-built mock scanner at least one day prior to scanning. Participants were presented with short movie clips of a children’s TV program during the acquisition of structural MRI.

#### 2.2.1 MRI data acquisition

We acquired four different types of MRI data for each participant: T1-weighed high-resolution anatomical images, T2*-weighed fMRI EPI, DWI, and GRASE MWI.

To co-register magnetic resonance images from different MRI sequence types as well as for probabilistic definition of the HC and the EVC ROIs based on gray-white matter segmentation and cortical parcellation, T1-weighed high-resolution anatomical images of the whole head were acquired with an MP-RAGE sequence and the following settings: TR = 8.10 ms, TE = 3.72 ms, flip angle = 8°, 220 slices, matrix size = 240 × 240, voxel size = 1 mm × 1 mm × 1 mm.

To define scene-selective regions of interest (ROIs), we obtained functional MRI during a scene localizer block design experiment using a blood oxygen level dependent (BOLD) sensitive T2*-weighted EPI sequence across 33 slices (TR = 2000 ms, TE = 30 ms, flip angle = 90°, FOV = 240 mm × 240 mm, voxel size = 3 mm × 3 mm × 3 mm, slice gap = 0.4 mm). Participants viewed a total of four 182 s (91 TRs) long runs. Each run comprised four blocks showing photographs of scenes, four blocks showing photographs of objects, and five rest condition blocks in which gray rectangles were shown. Each block contained 14 image presentations at 1 Hz. Stimuli were not repeated, except for one random occasion per block, which constituted a 1-back task to ensure sustained attention. Details of the scene localizer experimental design, stimuli, presentation and task are reported elsewhere (Meissner, Nordt, & Weigelt, 2019).

For fiber tracking and diffusion parameter analysis, a diffusion-weighted single-shot spin-echo EPI sequence along 33 isotropically distributed directions using a b-value of 1000 s/mm^2^ was obtained (TR = 7234 ms, TE = 89 ms, flip angle = 90°, 60 slices, matrix size = 128 × 128, voxel size = 2 × 2 × 2 mm). At the beginning of this sequence, one reference image was acquired without diffusion weighting (b = 0 s/mm^2^).

To examine the myelination state of white matter tracts, a 3D multi-echo (ME) gradient spin echo (GRASE) sequence with refocusing sweep angle was acquired (TR = 800 ms; TE = 10 - 320 ms, 32 echoes in steps of 10 ms, partial Fourier acquisition (z-direction: 50% overcontiguous slices, i.e. acquired slice thickness = 4 mm, reconstructed slice thickness =2 mm; y-direction: none), parallel imaging SENSE = 2.0, flip angle = 90°, 60 slices, matrix size = 112 × 112, voxel size = 2 × 2 × 2 mm, acquisition duration = 7.25 min).

#### 2.2.2 T1 anatomical image data processing and probabilistic ROI definition for EVC and HC

We excluded non-brain parts of the head in the T1 image using the FSL (FMRIB’s Software Library, RRID: SCR_002823, Jenkinson, Beckmann, Behrens, Woolrich, & Smith, 2012, version 6.0.3) BET tool (Smith, 2002).

To localize the EVC and the HC as additional ROIs in terms of key input/output areas of the scene-network ROIs, we used probabilistic atlases based on the T1-weighed images (in contrast to the scene-selective ROIs that are fMRI-based). Here, we used FreeSurfer (RRID: SCR_001847, version 6.0.0) for automated cortical parcellation, segmentation <u>and labelling</u> of the T1-weighted images. The details of the applied recon-all analysis pipeline have been described elsewhere (Dale, Fischl, & Sereno, 1999; Fischl et al., 2002; Fischl et al., 2004; Fischl, Sereno, & Dale, 1999; Ségonne et al., 2004) and the procedure has been shown to be valid for all age groups in our study to the same extent (Ghosh et al., 2010). To localize the EVC and the HC, we used FreeSurfer’s implemented probabilistic labels and atlases. For the EVC, Brodmann Area Maps labels for the primary and secondary visual area (V1/BA17 and V2/BA18) were used (labels “lh/rh.V1/V2_exvivo.thresh.label”, Fischl et al., 2008). For the HC, the hippocampus labels from the subcortical segmentation were used (aseg labels “lh/rh_hippocampus”).

We converted V1, V2, and HC FreeSurfer surface labels to ROI masks in FSL anatomical T1 space using FreeSurfer’s bbregister and mri_label2vol commands. Next, we registered V1, V2, and HC masks to DWI space using FSL FLIRT (FMRIB’s Linear Image Registration Tool, Greve & Fischl, 2009; Jenkinson, Bannister, Brady, & Smith, 2002; Jenkinson & Smith, 2001) for probabilistic tractography (Figure 1, top, middle). V1 and V2 were merged into a single EVC ROI, as later fiber tracking from V1 and V2 were barely distinguishable (see 2.2.4 DWI data processing, DTI model fitting, and probabilistic tractography).

**Figure 1:**
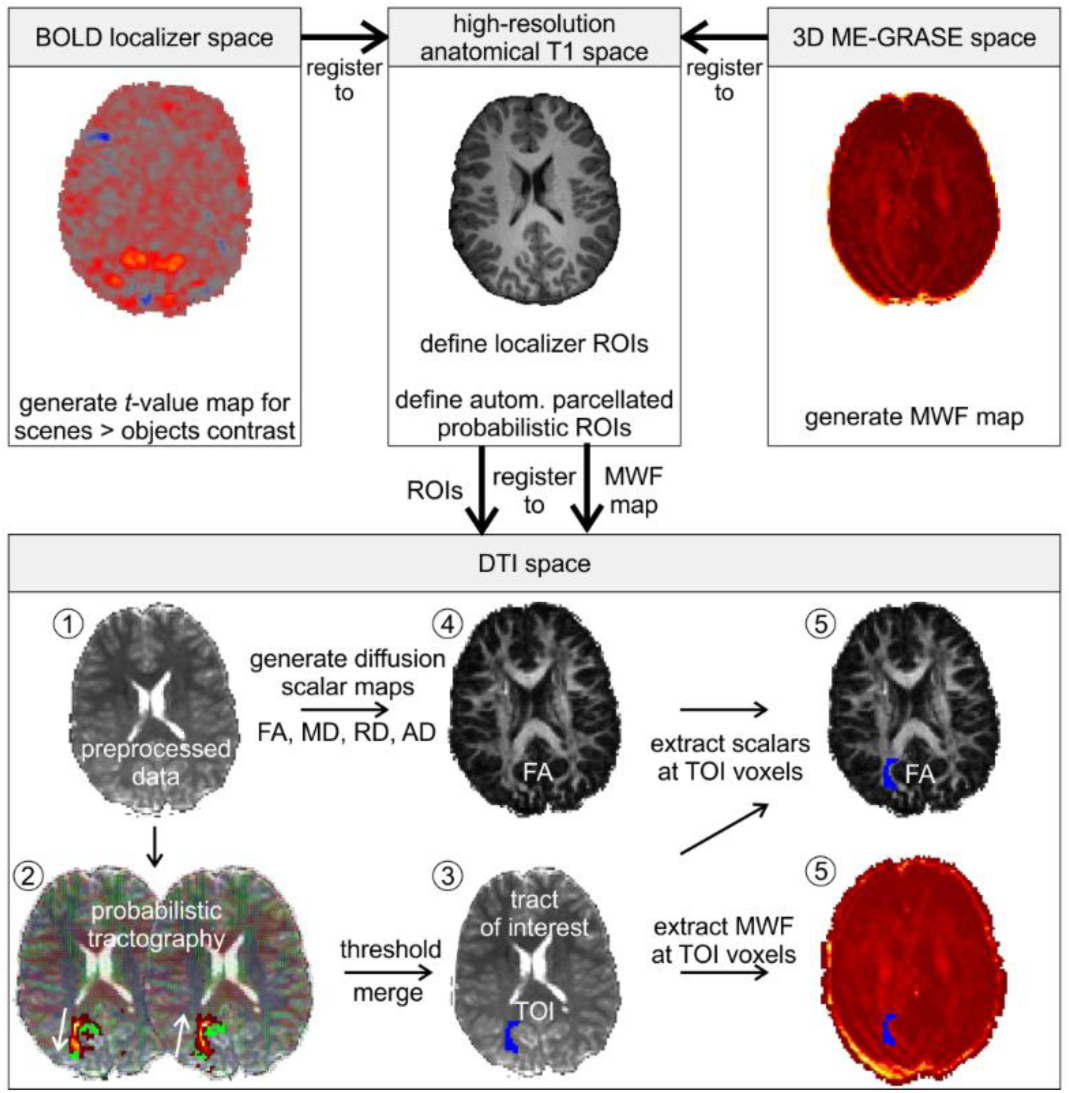
Analysis pipeline overview. The anatomical image in T1 space was used as a common intermediate registration template and connecting link between DTI space and non-diffusion spaces, i.e. BOLD and GRASE space. Numbers in circles indicate the order of analysis steps in DTI space. green = seed and target ROI (here: right RSC and OPA) for probabilistic tractography, heatmap arch = result for probabilistic tractography in which brighter colors indicate more samples crossing that voxel, blue = merged and thresholded TOI mask.

#### 2.2.3 fMRI data processing and functional ROI definition of PPA, RSC, and OPA

We used FSL FEAT (FMRI Expert Analysis Tool, FSL version 5.0.11, FEAT version 6.0.0, Woolrich, Ripley, Brady, & Smith, 2001) for preprocessing and statistical analysis of functional MRI localizer data. Preprocessing of functional data included brain extraction, slice time correction, motion correction, high-pass temporal filtering (cutoff: 91 s) and registration to the T1 anatomical image for each run in a first-level analysis. For each participant, first-level statistical results of all four runs were then entered into a mixed-effects second-level analysis (FLAME 1 option; Woolrich, Behrens, Beckmann, Jenkinson, & Smith, 2004), yielding a single statistical *t*-value maps for the scene > object contrast for each participant (Figure 1, top left).

To define scene-selective ROIs, we registered each participant’s *t*-value maps to her/his anatomical T1 image using sinc interpolation with FSL FLIRT (Figure 1, top left to top middle panel). Registration accuracy for each participant was visually examined and manually adjusted, if necessary. Using FSLeyes (version 0.22.6, McCarthy, 2018), we defined subject-specific PPA, RSC, and OPA in each hemisphere based on thresholded *t*-value maps (Figure 1, top middle; for exemplary scene-selective ROIs from multiple perspectives, see Supplementary Figure S1).

For the PPA and RSC, we included contiguous voxels whose scenes > objects contrast exceeded the *t*-value of 5.75. For the OPA, we chose a more liberal threshold of *t* > 4. This diverging and more liberal threshold was chosen because the OPA was rarely detectable at the original threshold of 5.75. Moreover, lowering the threshold for PPA and RSC to *t* > 4 in an attempt to use the same threshold for all ROIs resulted in large overlaps of PPA and RSC clusters, making a delineation impossible (cf. Meissner, Nordt, & Weigelt, 2019). Scene-selective ROIs did not overlap and only supra-threshold activation clusters at anatomically plausible locations as based on previous literature (Nasr et al., 2011; Weiner et al., 2018), but not putatively low-level driven clusters in the low-level visual cortices or other small distributed clusters were regarded. For all scene-selective ROIs, this approach yielded high detection rates that did not differ between age groups as determined using Fisher’s exact test (S1 Table). One 11-12yo was excluded from subsequent analyses, because activations for the scenes > objects contrast did not exceed the set thresholds.

For each participant, we registered the T1 anatomy to the b=0 DWI image using trilinear transformation with FSL FLIRT. Registration accuracy for each participant was visually examined and was found to be very accurate. No post-hoc manual adjustment was necessary for any participant. For a quality check of the cross-modal T1-to-DWI registration, see Supplementary Figure S2. The resulting transformation matrix was then used to register each scene-selective ROI from anatomical T1 space to DWI space using nearest neighbor interpolation (Figure 1, arrow from top middle to lower panel). For six ROIs in six participants, interpolation to the target space using the nearest neighbor algorithm failed. This was presumably due to their small size of 1 mm^3^ to 5 mm^3^ and the consequently difficult mapping to the quadrupled voxel size of 4 mm^3^ of the DWI target space. To still include these scene-selective ROIs in the analysis, we applied a trilinear interpolation approach in FSL FLIRT, which does not produce a binary mask, but a continuous probability map for that scene-selective ROI in target space. To obtain a binary scene-selective ROI mask again, we included all voxels at and above the probability map’s median value. This procedure was successful for all six scene-selective ROIs in which the original nearest neighbor algorithm failed.

#### 2.2.4 DWI data processing, DTI model fitting, and probabilistic tractography

For analysis of diffusion weighted data, we used FSL’s FDT (FMRIB’s Diffusion Toolbox, FSL version 6.0.3). Preprocessing of DWI data included eddy current and motion artefact correction using FSL eddy, diffusion gradient vectors reorientation to match the correction-induced rotations, as well as brain extraction (Figure 1, bottom, #1).

To evaluate white matter microstructural integrity in fiber tracts of interest, we fit diffusion tensors, modelled by three pairs of eigenvectors (*ε*_1,_ *ε* _2,_ *ε* _3_) and eigenvalues (λ_1,_ λ_2,_ λ_3_) that describe the direction and magnitude of water diffusion (unit: mm^2^/s × 10^−3^) along three orthogonal axes, to each voxel of our preprocessed DWI data using FSL DTIFIT. We then calculated axial, radial and mean diffusivity (AD = λ_1_, RD = (λ_1_ + λ_2_)/2, MD = (λ_1_ + λ_2_ + λ_3_)/3), as well as fractional anisotropy 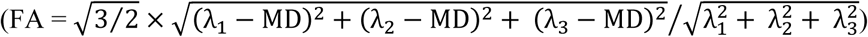 as diffusion parameters of interest (Figure 1, bottom, #4).

We performed probabilistic tractography on our data in native diffusion space using FSL BEDPOSTX and PROBTRACKX (Behrens et al., 2003; Behrens, Berg, Jbabdi, Rushworth, & Woolrich, 2007, FSL version 6.0.3) with the following settings: BEDPOSTX number of fibres per voxel = 2, ARD weight =1, burnin period =1000, number of jumps = 1250, sample every = 25, model = deconvolution with sticks, consider gradient nonlinearities = off. PROBTRACKX: loopcheck = on, onewaycondition = on, curvature threshold = 0.2, number of steps per sample = 2000, steplength in mm = 0.5, number of samples = 25000, volume fraction before subsidary fibre orientations are considered = 0.01, discards samples shorter than this threshold = 0.0, Sample random points within seed voxels = 4.0. For each participant, fiber tracking was done for 18 intrahemispheric tracts. In turn, each of the six scene-selective ROIs—as defined by our localizer (see 2.2.3 fMRI data processing and functional ROI definition of PPA, RSC, and OPA)—was set as the seed mask. For each seed mask, i.e. for each scene-selective ROI, four ipsilateral target masks were set: 1-2) the two other scene ROIs, 3) the HC, 4) the EVC (Figure 2). For these 18 seed-target pairs, probabilistic tractography was done in both directions. That is, after the initial seed-to-target tracking was done, a target-to-seed tracking estimated the same tract in reverse direction (Genç, Bergmann, Singer, & Kohler, 2011). For both directions, target masks were also set as waypoint and termination masks to ensure that only tracts would be retained that entered the target mask and that did not project onto other areas. Our rationale for employing this dual-direction approach was to control for any direction specific biases in probabilistic tractography, avoiding under-as well as overrepresentation of tract size or detectability. We refrained from interhemispheric tracking, as DWI and MWI parameters from these tracts would be masked by a major share of general corpus callosum development adding little—if any—insight into scene-network specific development (for corpus callosum development, see Meissner, Genç et al., 2019).

**Figure 2:**
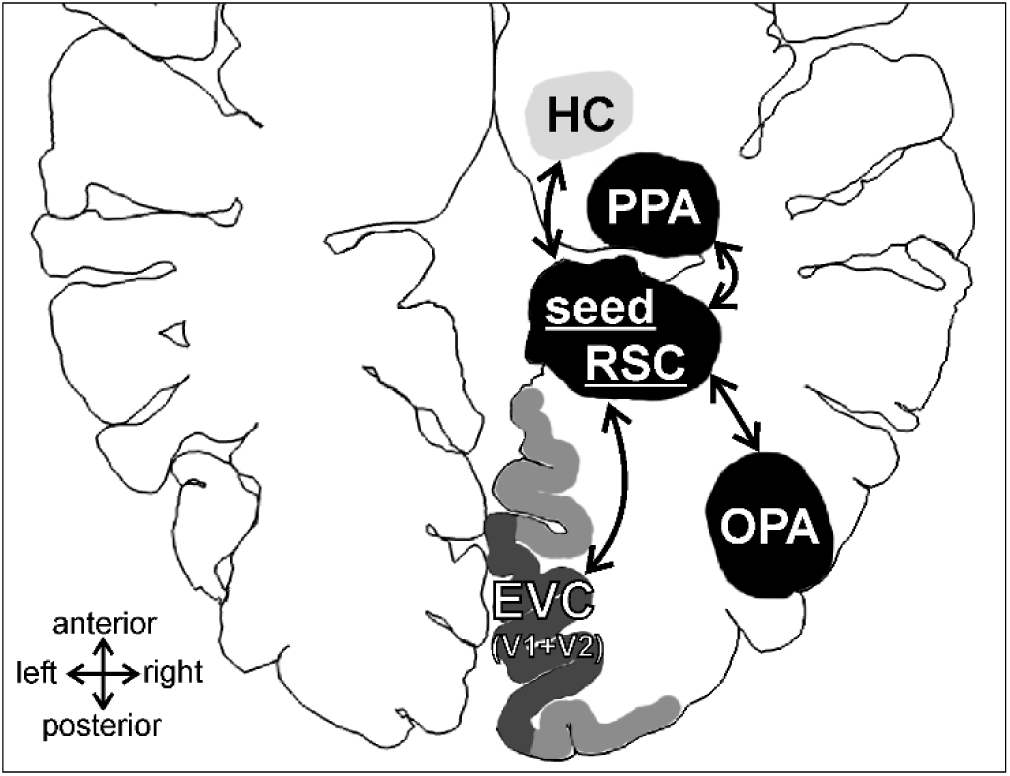
Schematic exemplary seed region (rRSC) with target regions and tract connections. For each scene-selective ROI, fiber tracking was done with the ipsilateral early visual cortex (EVC, i.e. a merged ROI of V1 and V2), hippocampus (HC), and the other two ipsilateral scene-selective ROIs (here: PPA and OPA). Note that in order to show all seed and target regions in one comprehensive display, this is a schematic drawing that does not reflect the appropriate absolute or relational position and size of target and seed regions.

For each voxel, the resulting probability maps indicate how many of the streamlines that successfully connected seed-to-target crossed this voxel. However, these probability maps include low-probability voxels that are likely to be spurious connections. To remove these spurious connections, we threshold individual tract maps at 20% of their robust maximum (99th percentile) value (Koldewyn et al., 2014; Yendiki et al., 2011) and then merged seed-to-target and target-to-seed tracts using a logical *and*-condition (Figure 1, bottom, #3; for exemplary tracts from multiple perspectives, see Supplementary Figure S1). Like other thresholding approaches, this accounts for systematically different ROI sizes. Moreover, in contrast to thresholding based on the number of initiated or successful streamlines, our approach provides a better interpretability, as the number of initiated or successful streamlines offers little insight into the actual probability map value distribution.

Originally, tracking from scene ROIs to the EVC was split into tracking to V1 and V2 based on their respective probabilistic FreeSurfer labels (2.2.2 T1 anatomical image data processing and probabilistic ROI definition for EVC and HC). However, visual inspection of final thresholded tracts revealed that tracts from scene ROIs to V1 and V2 were overlapping for the most part. Thus, we merged V1 and V2 ROIs to an EVC ROI and repeated the tracking procedure for the EVC ROI.

Weighted mean diffusion tensor imaging (DTI) metric values for each tract were obtained that considered each voxel’s DTI metric values and tract probability (Yendiki et al., 2011, Figure 1, bottom, #5). In detail, the weighed mean is the sum of all tract voxels’ DTI metric values multiplied with their tract probabilities. Each voxel’s tract probability is the crossing streamline count of the voxel divided by the sum of the crossing streamline count across all voxels.

#### 2.2.5 Myelin water fraction estimation

Parameter maps estimating the fraction of water molecules located between myelin layers— the myelin water fraction (MWF, MacKay et al., 1994)—for each voxel were created as described in Prasloski et al. (2012) (Figure 1, top right). MWF maps were then registered to native DWI space using FSL FLIRT. Here, for high-accuracy transformations, we employed a two-step procedure. First, we registered the TE = 10 ms image of the GRASE sequence to anatomical T1 space using trilinear transformation (Figure 1, arrow from top right to top middle panel). Registration accuracy for each participant was visually examined and manually adjusted, if necessary. The resulting transformation matrix was then used to register the MWF map to anatomical T1 space using sinc interpolation. Second, we registered the T1 anatomy to the b=0 DWI image using trilinear transformation (Figure 1, arrow from top middle to lower panel). The resulting transformation matrix was then used to register the MWF map from anatomical T1 space to DWI space using sinc interpolation. Weighted mean MWF values for each tract were obtained that considered each voxel’s MWF value and tract probability (in the same way as it was done for DWI, Figure 1, bottom, #5).

### 2.3 Neuroimaging data quality control

We screened preprocessed 4D DWI data and FA maps for visible artefacts that were not corrected by the preprocessing steps and excluded one 7-8yo participant from subsequent analyses. Further, to control for possible age group differences in DWI data quality, we quantified one registration-based and two intensity-based data quality measures implemented in the FSL eddy tool (Andersson, Graham, Zsoldos, & Sotiropoulos, 2016; Andersson & Sotiropoulos, 2016). For the registration-based measure, mean volume-to-volume voxel displacement in mm was analyzed (FSL eddy output: eddy_movement_rms). This measure is calculated by the eddy tool for each volume as the root mean square (RMS) of voxel displacements, was averaged across all volumes for each participant, and captures voxel displacements caused by both subject movement as well as eddy currents. With regard to subject movement, this measure is able to capture global, slow between-volume motion. Our analysis of variance (ANOVA) did not reveal a significant between-group difference (Figure 3 left: *F*(2,44) = 1.72, *p* = .842, η^2^ = .008)

**Figure 3:**
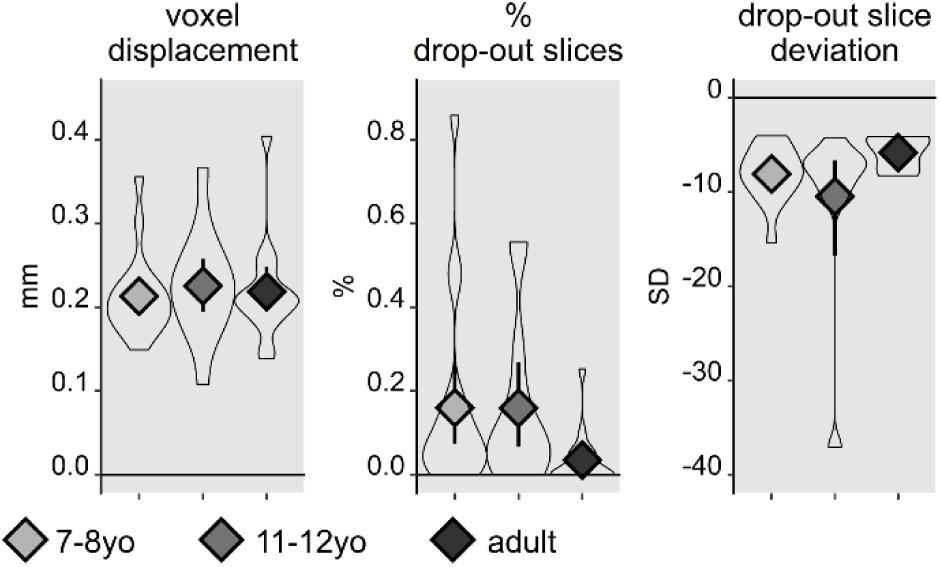
Between-group data quality comparison. Gray diamonds = group mean. Light gray = 7-8-year-old children, medium gray = 11-12-year-old children, dark gray = adults. Violin plots show the full distribution of the data. White circles = individual data points. Error bars show 95% confidence intervals for the mean.

For the intensity-based measures, we analyzed the percentage of slices with a significant signal intensity drop-out as implemented in the FSL eddy tool (-repol flag, Andersson et al., 2016). A significant signal intensity drop-out in a slice was defined as a mean voxel signal intensity that is more than 4 SD below the expected intensity (default setting for the -repol flag in FSL eddy version 6.0.3). This was to capture the effect of rapid within-volume motion (note: TR = 7234 ms). Our analysis of variance (ANOVA) did not reveal a significant between-group difference (Figure 3 middle: *F*(2,44) = 2.60, *p* = .085, η^2^ = .106) and the absolute percentage of drop out slices in all participants was below 1%. In addition, we analyzed the severity of signal dropout in the slices with signal intensity drop-out by comparing between age groups how many standard deviations (SD) the signal intensity of the dropout slices deviated from the mean difference between observed and expected signal intensity. Our analysis of variance (ANOVA) did not reveal a significant between-group difference (Figure 3 right: *F*(2,27) = 1.21, *p* = .315, η^2^ = .082). As the FSL eddy tool replaces slices with signal-drop out with and the age groups do not seem to differ in the number of replaces slices, the severity of the necessary corrections for signal intensity drop out or voxel displacements by subject motion or eddy currents, these factors are unlikely to contribute to any age group differences.

As the 3D signal acquisition method of the GRASE sequence is not volume-based, affine registration matrices and corresponding motion estimates, like for functional MRI or DWI cannot be computed for 3D ME-GRASE data. However, we visually screened all raw GRASE images as well as MWF maps for motion artefacts but found none.

### 2.4 Experimental design and statistical analysis

Our study investigated the effect of the between-subject factor age group (7-8yo, 11-12yo, adults) on the outcome variables MWF, FA, MD, RD, and AD for six scene-selective fiber tracts. In an exploratory analysis, further tracts were tested between all six scene-selective ROIs and the EVC and HC, respectively. To test for differences between age groups, we employed analysis of variances (ANOVAs) for each fiber tract independently. As multiple statistical comparisons of the six tracts might cause an inflation of α-error, we used the Bonferroni method to adjust the default significance threshold of α = .05 to α = .0083 (0.05/6 tracts). To improve the usability of our results for colleagues whose research interest focuses on one or a particular region or tract of interest only, we also report age group effects that reached the uncorrected significance threshold of α = .05 in the results section. Statistical data analysis was performed using R (version 3.6.0, RRID: SCR_001905, R Core Team, 2019) in RStudio (version 1.2.1335; RRID: SCR_000432). No part of the study procedures, experimental design, or statistical analyses were pre-registered prior to the research being conducted.

## 3 Results

This study combined myelin water imaging with a functional MRI scene localizer and DWI-based tractography to determine the degree of myelination in white matter tracts underlying the cortical scene-network in three age groups. We examined possible differences in myelin water fraction between eighteen 7-8yo, thirteen 11-12yo, and sixteen adults to examine if the scene-network’s white matter structural connectivity follows a similar or divergent pattern in reference to scene-network’s functional development. Further, we investigated connections between the scene-network and strongly connected areas, such as EVC and the HC. In an extended analysis, we repeated our analyses for DTI parameters to relate DTI and MWF results patterns.

### 3.1 Myelin water imaging

Regarding within-scene-network tracts, the MWF in fibers connecting the right PPA and OPA increased with age (*F*(2,22) = 6.02, *p* = .0082, η^2^ = .354, Figure 4). We observed further increases, albeit not surviving Bonferroni correction, for the left and the right RSC-OPA tract (left: *F*(2,22) = 3.64, *p* = .0432, η^2^ = .248; right: *F*(2,26) = 4.59, *p* = .0196, η^2^ = .261).

**Figure 4:**
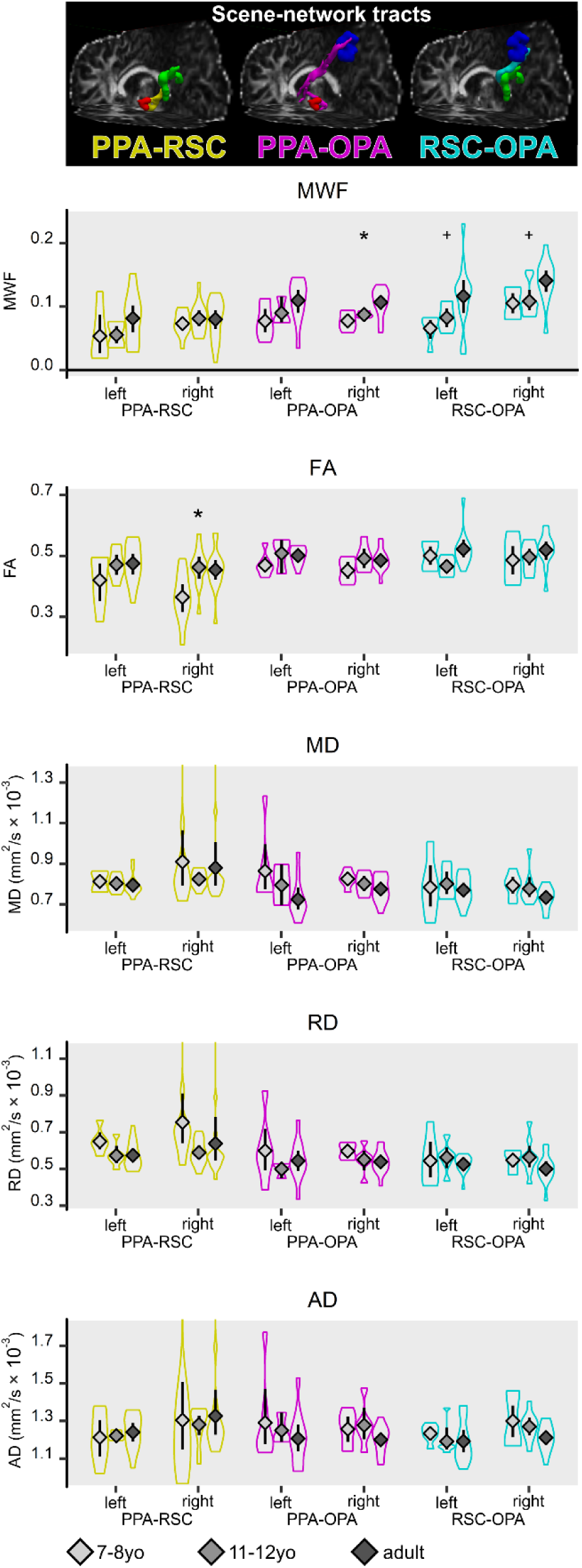
Myelin water fraction (MWF) and DTI parameters for intrahemispheric connections between scene-network areas. Gray diamonds = group mean. Light gray = 7-8-year-old children, medium gray = 11-12-year-old children, dark gray = adults. Error bars show 95% confidence intervals for the mean. Violin plots show the full distribution of the data. Asterisks and plus signs indicate significance with Bonferroni correction (α = .0083) and without correction for multiple comparisons (α = .05), respectively.

For connections between the HC and scene-network areas as well as for connections between the EVC and scene-network areas we did not find any age group differences in MWF that survived Bonferroni-correction (Figure 5, first row). Non-significant increases, i.e. not surviving Bonferroni-correction, were observed for both HC connections and EVC connections (left RSC-HC: *F*(2,38) = 4.31, *p* = .0205, η^2^ = .185; left OPA-HC: *F*(2,36) = 5.27, *p* = .0098, η^2^ = .227; right OPA-HC: *F*(2,34) = 5.28, *p* = .0101, η^2^ = .237; left RSC-EVC: *F*(2,38) = 4.34, *p* = .0200, η^2^ = .186; right OPA-EVC: *F*(2,35) = 3.612, *p* = .0375, η^2^ = .171; Figure 5, first row)

**Figure 5:**
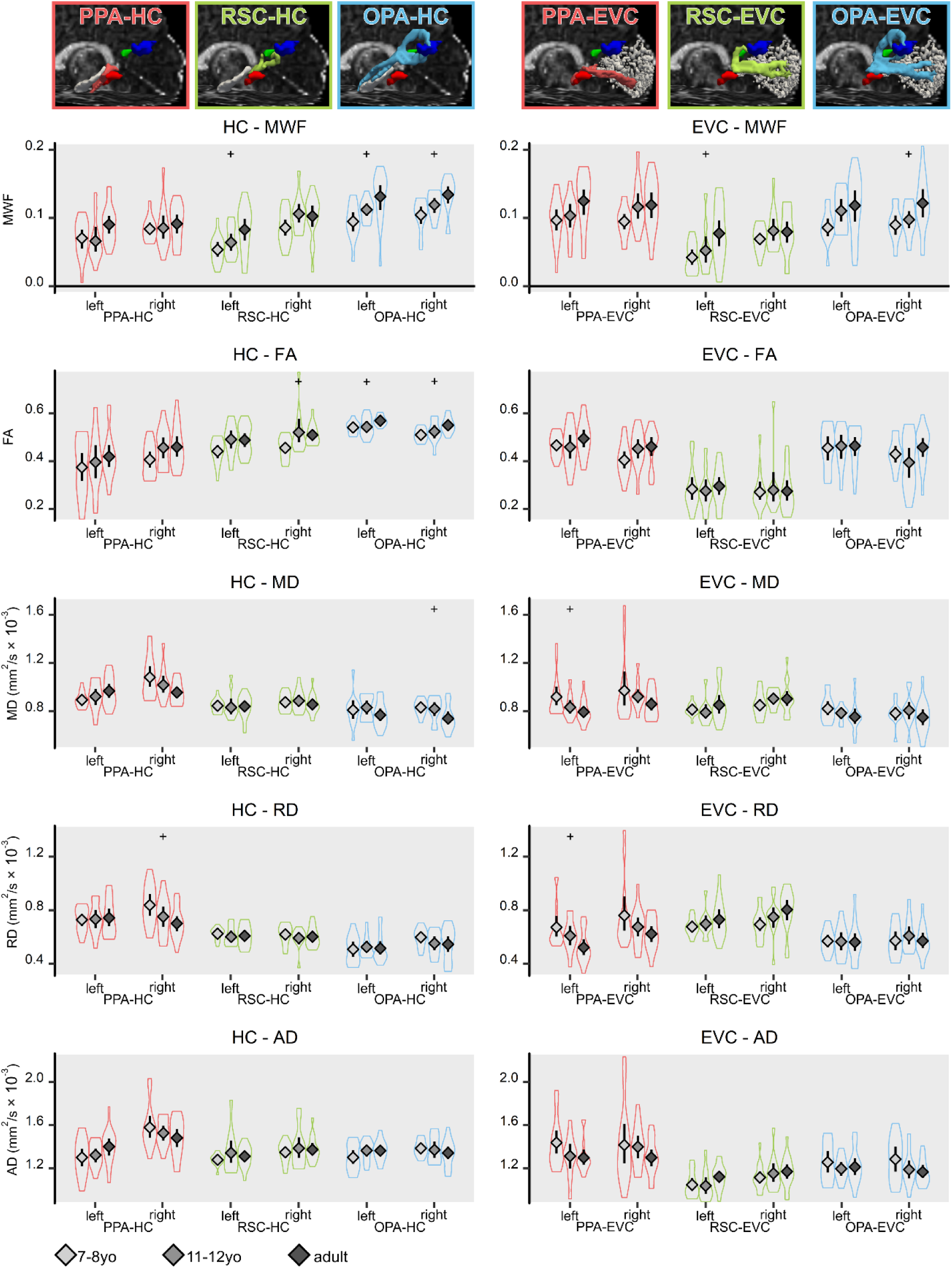
Myelin water fraction (MWF) and DTI parameters for connections between scene-network areas and the hippocampus (HC, left), early visual cortex (EVC, right). Gray diamonds = group mean. Light gray = 7-8-year-old children, medium gray = 11-12-year-old children, dark gray = adults. Error bars show 95% confidence intervals for the mean. Violin plots show the full distribution of the data. Plus signs indicate non-significant trends that did not meet our with Bonferroni correction threshold of α = .0083, but only the conventional threshold of α = .05.

### 3.2 Extended analysis of DTI parameters

Regarding within-scene-network tracts, the age group difference in the right PPA-OPA tract observed for MWF was not mirrored in DTI parameters. For FA, we found increasing values with age in the right PPA-RSC tract, (*F*(2,36) = 6.76, *p* = .0032, η^2^ = .273, Figure 4, second row). No other tract showed age effects for FA. Concerning the other DTI parameters—MD, RD, and AD—we did not find any tracts with age group differences (Figure 4, third, fourth, and fifth row).

As for MWF, we did not find any tracts with age group differences that passed the Bonferroni-correction for HC-scene-network connections or EVC-scene-network connection (Figure 5, second, third, fourth, and fifth row). For FA, non-significant increases were observed in right RSC-HC (*F*(2,40) = 4.60, *p* = .0159, η^2^ = .187), left OPA-HC (*F*(2,36) = 3.97, *p* = .028, η^2^ = .181), and right OPA-HC tracts (*F*(2,34) = 4.38, *p* = .020, η^2^ = .205) (Figure 5, second row). For MD, non-significant increases were observed in right OPA-HC tracts (*F*(2,34) = 3.96, *p* = .0284, η^2^ = .189) and left PPA-EVC tracts (*F*(2,40) = 3.68, *p* = .0342, η^2^ = .155) (Figure 5, third row). For RD, non-significant increases were found in right PPA-HC tracts (*F*(2,40) = 3.72, *p* = .0331, η^2^ = .157) and left PPA-EVC tracts (*F*(2,40) = 4.57, *p* = .0164, η^2^ = .186) (Figure 5 fourth row). For AD no non-significant increases were found (Figure 5 fifth row).

### 3.3 Control analyses

The volumes of PPA and OPA, but not RSC, differed between age groups, with larger scene-selective ROIs in older age groups (cf. Meissner, Nordt, & Weigelt, 2019; volume was assessed in DTI space). We did not find age group differences for volume of the EVC or the HC. As seed- or target-ROI size could potentially influence tract size, we controlled for the possibility that any of the observed age effects in MWI or DTI metrics were confounded by age-related tract volume differences. We compared tract volume between age groups using ANOVAs and found differences in the right EVC-PPA tract (*F*(2,40) = 6.10, *p* = .0049, η^2^ = .234; which did not show any MWF or DTI age group differences), but not in any other tracts. Thus, age group differences in MWF or DTI metrics are unlikely to stem from tract size differences between age groups.

The nine tracts in each participants’ hemisphere were likely to show some degree of overlap. That is, some voxels’ MWF and DTI metrics would influence two or more tracts. In general, we do not deem these overlaps as problematic. We strived to investigate the maturity along the whole length of each tract and microstructural changes in overlap regions, i.e. in shared white matter pathways, would naturally contribute to both tract’s signal transmission. Still, to inform about the level of overlap, we calculated the mean overlap across all participants for each scene-network tract with the other two scene-network tracts in the respective hemisphere. The largest overlap between scene-network tracts was evident between the left RSC-OPA and PPA-OPA tract (the overlap was 14.26% of RSC-OPA’s tract size and 10.71% of PPA-OPA’s tract size). The second largest overlap between scene-network tracts was evident between the left PPA-RSC and PPA-OPA tract (the overlap was 10.42% of PPA-RSC’s tract size and 5.46% of PPA-OPA’s tract size). All other overlaps of scene-network tracts were below 10% of any of the two overlapping tracts. Concerning the scene-network-HC/EVC tracts, we also calculated the mean overlap across all participants for each scene-network-HC/EVC tract with all scene-network tracts in the respective hemisphere to find out how much of the scene-network-HC/EVC tracts was already “covered” by the scene-network tracts. The largest overlap was evident for the RSC-HC tract, which shared 32.44 % of its voxels with the PPA-RSC tract in the left hemisphere and 23.71% of its voxels with the PPA-RSC tract in the right hemisphere. For the ten other tracts, between 9.70% and 20.02% of their voxels were shared with other tracts.

## 4 Discussion

Myelin emergence and further maturation is a crucial step in brain development (Flechsig, 1920). While myelin development trajectories for white matter beyond late childhood are still unclear, it is established that the rate of change and the point at which an adult level is reached is region specific (Yakovlev & Lecours, 1967). Further, myelin maturation was shown to interact with functional organization and behavior (e.g. Bengtsson et al., 2005; Yeatman, Dougherty, Ben-Shachar, & Wandell, 2012). Here, we compared MWF in white matter tracts underlying the visual scene-network between 7-8yo, 11-12yo and adults. We found increasing MWF in the right PPA-OPA tract but not in any other tracts that connect scene-network regions with each other or with the HC or EVC (for non-significant trends, see section 3 Results).

### 4.1 Connections between scene-network areas

Our findings provide evidence for a protracted development of white matter tract that connect the scene-network regions PPA and OPA. Integrating our results with recent findings of functional cortical development in scene-selective areas during and beyond childhood opens up the possibility of structure-function interactions, i.e. influences of structural development on functional development, or vice versa. Previous studies established that the RSC is adult-like in middle childhood already (Jiang et al., 2014; Meissner, Nordt, & Weigelt, 2019), but that the PPA and the OPA show prolonged development in terms of cluster size, and scene-selectivity (Golarai et al., 2007; Meissner, Nordt, & Weigelt, 2019). This correspondence could imply that if structure-function interactions exist, they need the involvement, i.e. development, of both cortical ends of a tract, as no tracts to the RSC were found to show significant age group differences. Alternatively, no interactions might exist or the involvement of just one cortical end of a tract is sufficient. That is, structural and functional development might be independent and the correspondence of our structural findings with previous functional findings might just be a coincidence, possibly indicated by several non-significant trends in tracts involving the RSC. However, as cortical structure, function, and associated cognitive abilities have been associated with white matter structural development (Fields, 2008; Gomez et al., 2017), we speculate that a completely independent development of structure and function is unlikely.

While we do not report behavioral measures of scene processing in this study, previous studies have shown that the behavioral processing of scenes follows a protracted trajectory. This opens up the possibility that the PPA-OPA tract development observed in our study might be responsible for some aspects of behavioral development regarding the processing of scenes. For example, children younger than 6 years lack the ability to scan task-appropriate and informative parts of scenes. Further, the duration of fixations and the amplitude of saccades in scene exploration reaches adult levels around 10 years only (Fioravanti, Inchingolo, Pensiero, & Spanio, 1995; Helo, Pannasch, Sirri, & Rämä, 2014; Mackworth & Bruner, 1970; Vurpillot, 1968). Perceptual discrimination and change detection of scenes as well as recognition memory for scenes reaches adult levels between 10 years of age and adolescence (Chai et al., 2010; Dirks & Neisser, 1977; Fandakova, Leckey, Driver, Bunge, & Ghetti, 2019; Hock, Romanski, Galie, & Williams, 1978; Meissner, Nordt, & Weigelt, 2019; Weigelt et al., 2014). A more efficient connection between PPA, with its spatial layout and context-processing properties and OPA, with its first-person perspective spatial relations-processing properties could contribute to these observable developments.

Recent findings indicate that myelin in a majority of major white matter tracts increases from childhood into young adulthood (Meissner, Genç et al., 2019). In this study, major tracts that may partly subserve scene-network connecting tracts—i.e. the inferior longitudinal fasciculus—showed moderate myelin increases from middle childhood to adulthood. However, this remains speculative until other studies can replicate these findings, as our study’s population was equal to the one investigated in Meissner, Genç et al. (2019). Given that only one scene-network tract showed significant age group differences, it seems that, in contrast to major long distance tracts, short-distant specifically defined scene-network tracts do not necessarily follow the same myelination pattern. However, future higher-powered work with larger sample sizes might elucidate these associations further by confirming if age group differences are really specific to the PPA-OPA tract or a more general phenomenon within the scene-network and thus resembling the developmental trajectory of major long white matter tracts, as well as cortical gray matter myelin content (Carey et al., 2018; Grydeland, Walhovd, Tamnes, Westlye, & Fjell, 2013; D. J. Miller et al., 2012; Shafee, Buckner, & Fischl, 2015). For now, our findings suggest that short-distance modality-specific white matter tracts are independent of long-distance white matter development or general gray matter myelin development and that myelin increases in scene-network white matter are tract specific.

### 4.2 Connections between the scene-network, hippocampus, and EVC

Connections from scene-network areas to the HC or the EVC did not indicate increasing myelination from middle childhood to adulthood. This suggests that white matter connections from HC and EVC are generally mature at an earlier time that within-scene-network connection. For the EVC this agrees with evidence of early maturity of cortical structures and most visual functions (Braddick & Atkinson, 2011). In contrast, memory and context-related cognitions supported by the hippocampus were shown to reach adult level in adolescence only (for scene specific effects, see e.g. Meissner, Nordt, & Weigelt, 2019; Weigelt et al., 2014). In summary, our findings suggest that white matter connections that connect the scene-network to major input-/output areas are unlikely to show drastic developments and therefore unable to explain age-related changes in scene-related cognition.

### 4.3 Diffusion tensor imaging interpretation

None of the investigated DTI parameters (FA, MD, RD, AD) mirrored our MWI findings of age group differences in the right PPA-OPA tract. This missing correspondence might be explained by the fact that fiber geometry has a particularly high influence on DTI parameters in small tracts—like connections between scene-network areas. This high influence is due to a higher probability that two tracts with diverging principal diffusion directions cross, branch, or merge within one voxel (Feldman et al., 2010). Thus, especially in small tracts, the use of DTI parameters as a proxy for myelin is problematic and might not reflect myelination but rather other microstructural changes, such as fiber geometry (Friedrich et al., 2020; Moura et al., 2016). For example, a recent study on the same study population that investigated major large tracts found a comparatively higher correspondence between DTI and MWF effects (Meissner, Genç et al., 2019).

### 4.4 Limitations

With our investigation on the scene-network white matter development our study provides an important contribution to an integrated understanding of how the scene-network of PPA, RSC, and OPA develops. However, the methodological approach of our study has certain limitations that are discussed below.

Previous studies indicate that the optimal ratio of low-b acquisitions and high-b acquisitions is 0.1 (Jones, Horsfield, & Simmons, 1999) to 0.2 (Alexander & Barker, 2005). Consequently, an optimal ratio for our protocol of 33 high-b acquisitions, would be achieved with 3-6 low-b (b=0) acquisitions (Mukherjee, Chung, Berman, Hess, & Henry, 2008). However, software limitations (Mukherjee et al., 2008) at the time of the recordings prevented the acquisition of more than one b=0 for a DWI session. Future studies with optimal scan protocols should therefore test whether our results are replicable.

The majority of studies in cognitive and developmental cognitive neuroscience quantify fiber geometry and microstructural properties of fiber tracts by means of the tensor model. This model assumes that, in each voxel, there is a unique orientation of fibers, the direction of which is represented by the tensor’s main eigenvector (Mori & Tournier, 2014). However, large portions of white matter voxels contain multiple fiber orientations (Jeurissen, Leemans, Tournier, Jones, & Sijbers, 2013). Therefore, tensor models are naturally limited to deal with voxels containing multiple fiber orientations (e.g. crossing fibers). Non-tensor-based models such as the method of constrained spherical deconvolution (CSD, Tournier, Calamante, Gadian, & Connelly, 2004) can be used to estimate the distribution of fiber orientations present within each voxel. With this method, the signal is measured by means of a high angular resolution diffusion imaging (HARDI) session that —in short— lead to a robust determination of the fiber orientations in voxel and have been shown to be superior to DWI-based tractography in some contexts (Farquharson et al., 2013). However, our scan protocol is not optimized for these kinds of methods, so future should test whether the development of scene-network-specific white matter tracts show similar trajectories if CSD-fiber tracking is employed.

As stated above, large portions of white matter voxels contain multiple fiber orientations (Jeurissen et al., 2013). While MWF is more specific to myelin than DTI-derived parameters, it is still not clear which exact axons within a voxel contribute to the MWF. Pathways that overlap and cross our scene-specific pathways in a substantially different direction, i.e. axon populations within voxels of our scene-pathways that do not serve the scene-specific connection, could influence the MWF at these points. Upon visual inspection, we could not identify major pathways that cross the scene-specific pathways on a regular basis except for the superior longitudinal fasciculus (SLF) that crosses the PPA-OPA tract. Aside from the fact that only for 1/6 of the tracts, a potential crossing tract exists that is reliably and automatically traceable with our data, even for the SLF and the PPA-OPA tract, the actual overlap was minimal: Only two participants showed an overlap of more than 10% shared voxels between the SLF and the PPA-OPA tract (each in only one hemisphere) and the maximum overlap was 16.4% shared voxels. Thus, the possible bias or the possibility to identify a bias by quantifying MWF in cross-over and non-cross-over voxels is very small.

MWF has shown strong qualitative and quantitative correspondence with histological markers for myelin (Laule et al., 2006; Laule et al., 2008). Still, it is important to be aware of potential confounding factors that may influence in vivo measurement of myelin water—which remains an indirect measure (MacKay & Laule, 2016). The most important factor is movement of water from myelin bilayers during the measurement. The T2 decay curve approach, which is used for the MWF, assumes that water molecules stay in the myelin bilayers for long times compared to the decay curve measurement time. However, at least studies in rodent spines indicate that during the measurement water molecules might be able to move from myelin in sufficiently fast rates to cause artificially low MWF (Harkins, Dula, & Does, 2012; Levesque & Pike, 2009). At the same time, other studies in animals indicate that water exchange does not play a considerable role in MWF measurements (Stanisz, Kecojevic, Bronskill, & Henkelman, 1999). The effect of water exchange in humans has not yet been accurately quantified; however, it seems likely that measured MWF are slight underestimates of the true MWF (Kalantari, Laule, Bjarnason, Vavasour, & MacKay, 2011). Moreover, future studies should compare our findings to alternative in vivo MRI measures that are able to quantify myelin architecture, such as magnetization transfer imaging (MacKay & Laule, 2016), bound pool fraction (Stikov et al., 2011), or myelin density (Sepehrband et al., 2015).

### 4.5 Outlook

Here, we investigated the development of myelin in white matter tracts subserving the cortical visual scene-network for the first time. We established that myelin seems to increase in at least one within-scene-network tract. This result is exciting in so far as it demonstrates that the protracted scene-network development between childhood and adulthood is not limited to functional changes (Meissner, Nordt, & Weigelt, 2019), but also includes maturation of underlying structures that are not directly part of the cortical network. We are positive that our study opens up two further directions going forward. First, our cross-sectional study paves the way for large-scale longitudinal studies with short time intervals over an extended period of time and a high number of participants that combine behavioral testing, fMRI, DWI-tractography, and MWI, which could tap into the important question of structure-function-development in more detail. Second, next to the scene-network, other cortical category-specific high-level vision areas form networks. For example, face processing is supported by a core network with modules in the fusiform gyrus, inferior occipital gyrus, and superior temporal sulcus, for which evidence also suggests a prolonged functional development (e.g. Golarai et al., 2007; Nordt, Semmelmann, Genç, & Weigelt, 2018, for a review see Haist & Anzures, 2017). Only little evidence, based on DTI analysis of major white matter tracts, exists that hints at possible emerging structure-function relations in the developing face processing system (Scherf, Thomas, Doyle, & Behrmann, 2014). Using MWI and tractography of individual, short-range, face-area-specific tracts, future research might corroborate these first findings and shed more light on structural white matter development as a contributing developmental factor on the long way to (face) perception expertise.

## Acknowledgements

We thank our team of student assistants and interns for assisting in stimulus creation, pilot testing, subject recruiting, and data collection. We acknowledge the support of the Neuroimaging Centre of the Research Department of Neuroscience at Ruhr University Bochum’s teaching hospital Bergmannsheil and Philips GmbH, Germany. We thank all participants and their parents for participating in this study.

## Data and code availability statement

All code used for data analysis, all tasks and stimuli, as well as anonymized raw data are openly available at the Open Science Framework (https://doi.org/10.17605/osf.io/dbg83) with one exception. For legal reasons (shared intellectual property with previous collaborators that did not author this manuscript and do not consent to open publication), the MWF parameter map generating algorithm cannot be shared openly. However, it is available upon request via e-mail to BM if the intended use is for academic non-commercial purposes. In addition to the possibility of requesting the algorithm to create the MWF maps, we have also uploaded all MWF maps to the Open Science Framework repository.

## Transparency statement

We report how we determined our sample size (2.1 Participants), all data exclusions (2.1 Participants), all inclusion/ exclusion criteria (2.1 Participants), whether inclusion/exclusion criteria were established prior to data analysis (2.1 Participants), all manipulations (2.4 Experimental design and statistical analysis), and all measures in the study (2.2 Neuroimaging).

## Ethics statement

The Ruhr University Bochum Faculty of Psychology ethics board approved the study (proposal no. 280). All participants as well as children’s parents gave informed written consent to participate voluntarily.

## CRediT author statement

**Tobias W. Meissner**: Conceptualization, Formal Analysis, Investigation, Data Curation, Writing—Original Draft, Writing—Review and Editing, Visualization, Project Administration, Funding Acquisition; **Erhan Genç**: Methodology, Formal Analysis, Resources, Writing—Review and Editing, Supervision; **Burkhard Mädler**: Methodology, Writing—Review and Editing; **Sarah Weigelt**: Conceptualization, Resources, Writing—Review and Editing, Supervision, Funding Acquisition

