## Supplementary Material for "Myelin development in visual scene-network tracts beyond late childhood: A multimethod neuroimaging study"


### S1 Table

S1 Table: Number and percentage of detected ROIs

| ROI | 7-8yo (n = 18) | 11-12yo (n = 13) | adults (n = 16) | test statistic | | *p*-value |
|---|---|---|---|---|---|---|
| lPPA | 17, 94.44 % | 11, 84.62 % | 16, 100 % | $\chi^2_{(2)}$ = | 2.87 | *p* = .266 |
| rPPA | 15, 83.33 % | 13, 100 % | 16, 100 % | $\chi^2_{(2)}$ = | 5.16 | *p* = .102 |
| lRSC | 15, 83.33 % | 11, 84.62 % | 16, 100 % | $\chi^2_{(2)}$ = | 2.90 | *p* = .241 |
| rRSC | 16, 88.89 % | 13, 100 % | 16, 100 % | $\chi^2_{(2)}$ = | 3.37 | *p* = .186 |
| lOPA | 13, 72.22 % | 11, 84.62 % | 16, 100 % | $\chi^2_{(2)}$ = | 5.16 | *p* = .076 |
| rOPA | 14, 77.78 % | 12, 92.31 % | 15, 93.75 % | $\chi^2_{(2)}$ = | 2.36 | *p* = .411 |

### S1 Figure

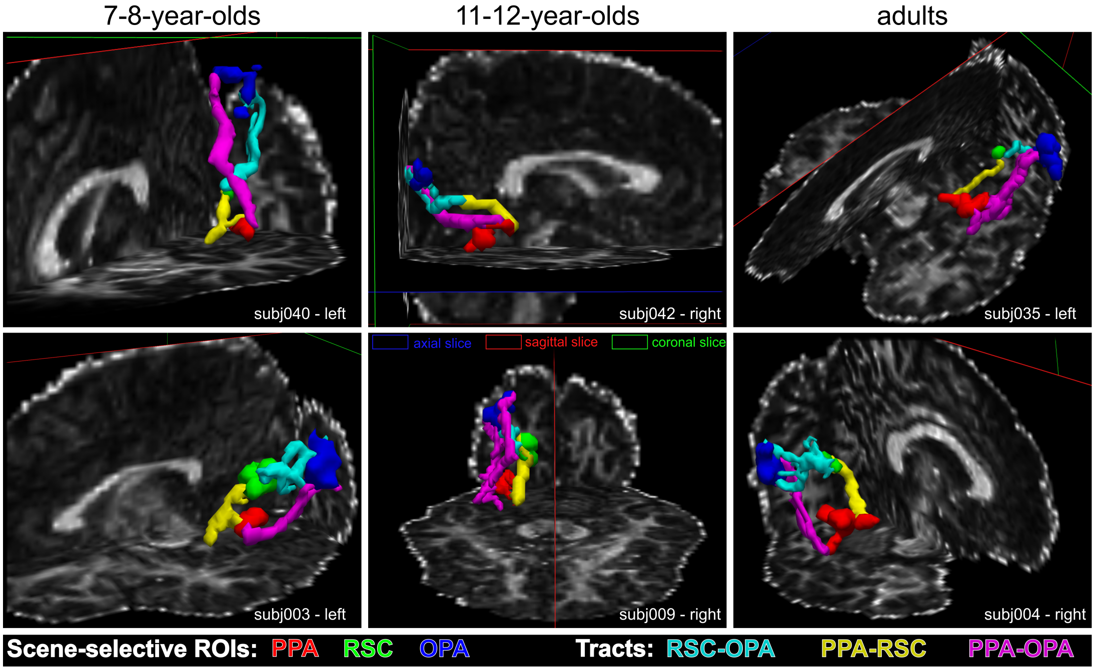


S1 Figure: Exemplary scene-selective ROIs (red = PPA, green = RSC, blue = OPA) and connecting fiber tracts (cyan = RSC-OPA, yellow = PPA-RSC, magenta = PPA-OPA) in two 7-8-year olds, two 11-12-year olds, and two adults shown from multiple different perspectives. ROIs and tracts are reconstructed in native 3D DTI space on top of a FA map. ROIs, tracts, and FA map are smoothed for better visualization. Tracts are displayed in uniform color for easier identification; for an example of a tract probability (heat-)map, see Figure 1, bottom, left.

### S2 Figure

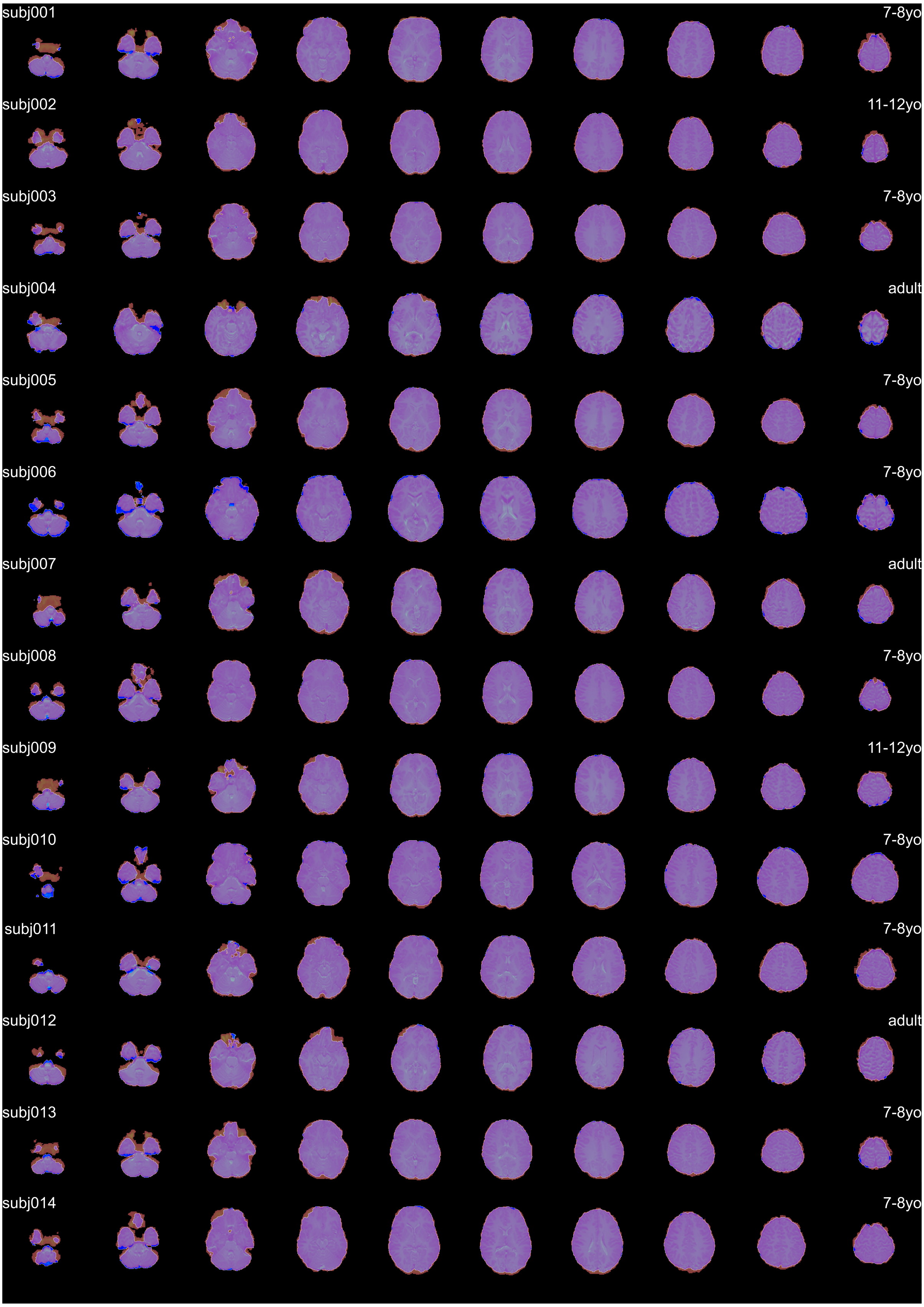
subj001
7-8yo
subj002
11-12yo
subj003
7-8yo
subj004
adult
subj005
7-8yo
subj006
7-8yo
subj007
adult
subj008
7-8yo
subj009
11-12yo
subj010
7-8yo
subj011
7-8yo
subj012
adult
subj013
7-8yo
subj014
7-8yo

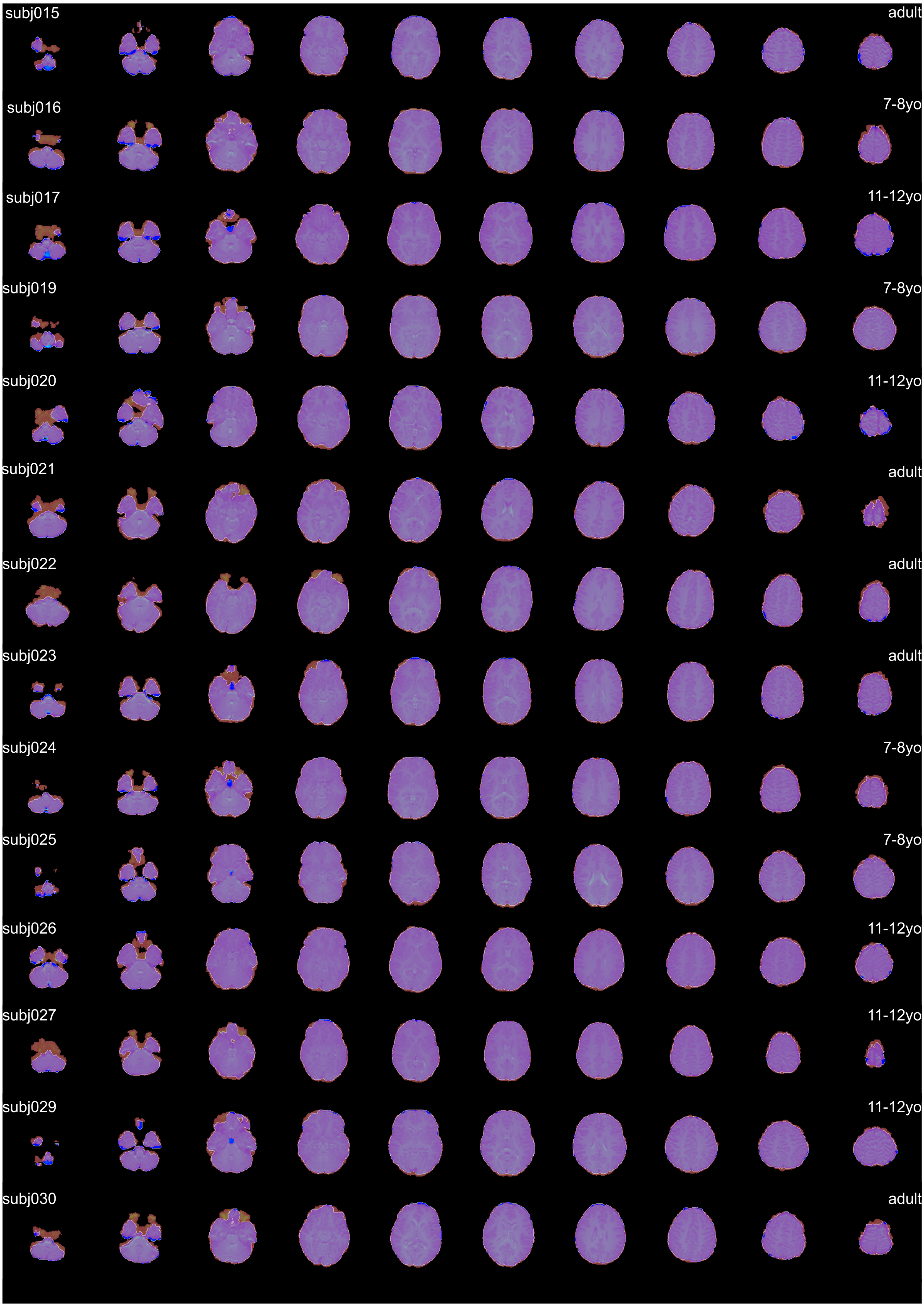
subj015
adult
subj016
7-8yo
subj017
11-12yo
subj019
7-8yo
subj020
11-12yo
subj021
adult
subj022
adult
subj023
adult
subj024
7-8yo
subj025
7-8yo
subj026
11-12yo
subj027
11-12yo
subj029
11-12yo
subj030
adult

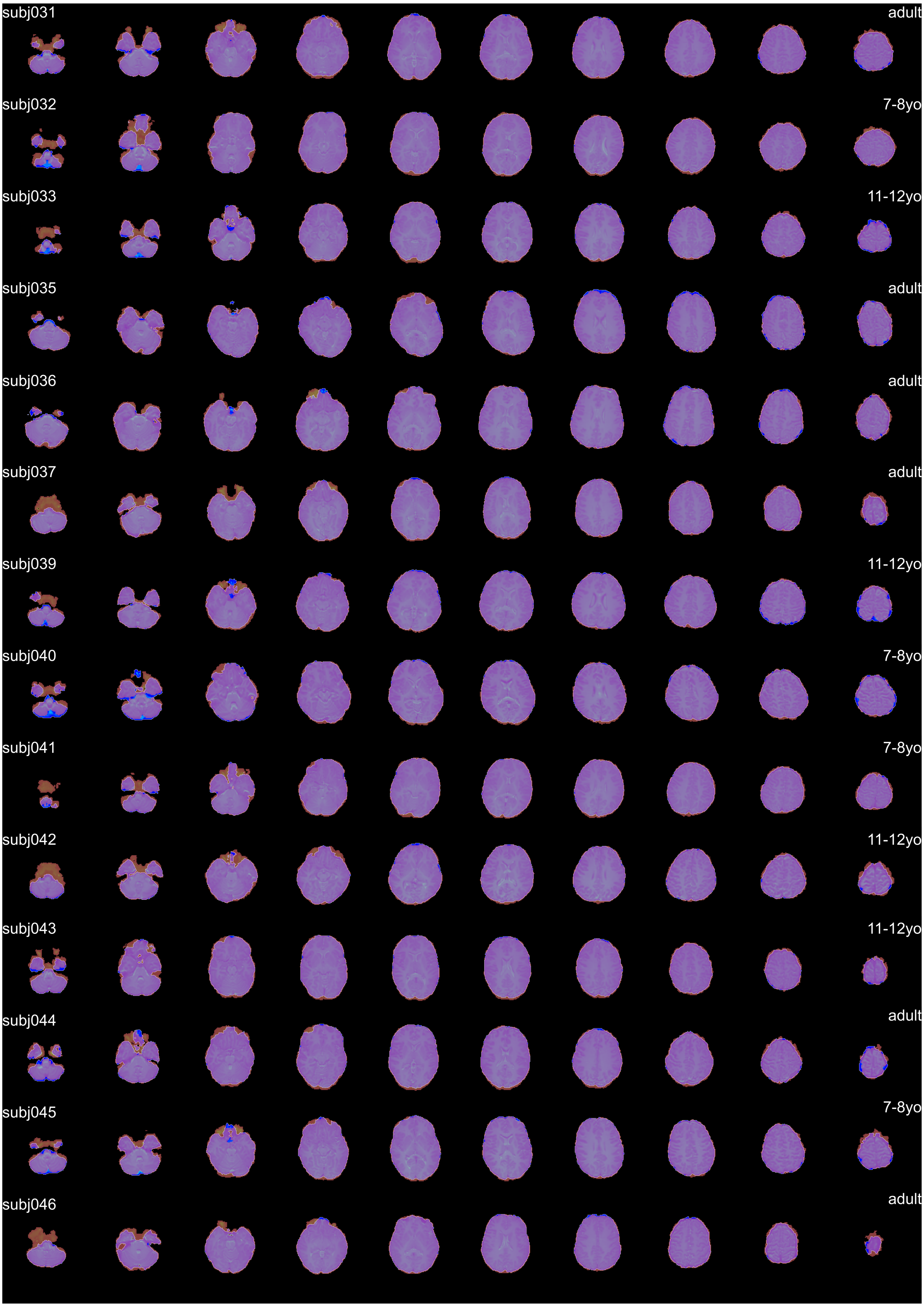
subj031
adult
subj032
7-8yo
subj033
11-12yo
subj035
adult
subj036
adult
subj037
adult
subj039
11-12yo
subj040
7-8yo
subj041
7-8yo
subj042
11-12yo
subj043
11-12yo
subj044
adult
subj045
7-8yo
subj046
adult

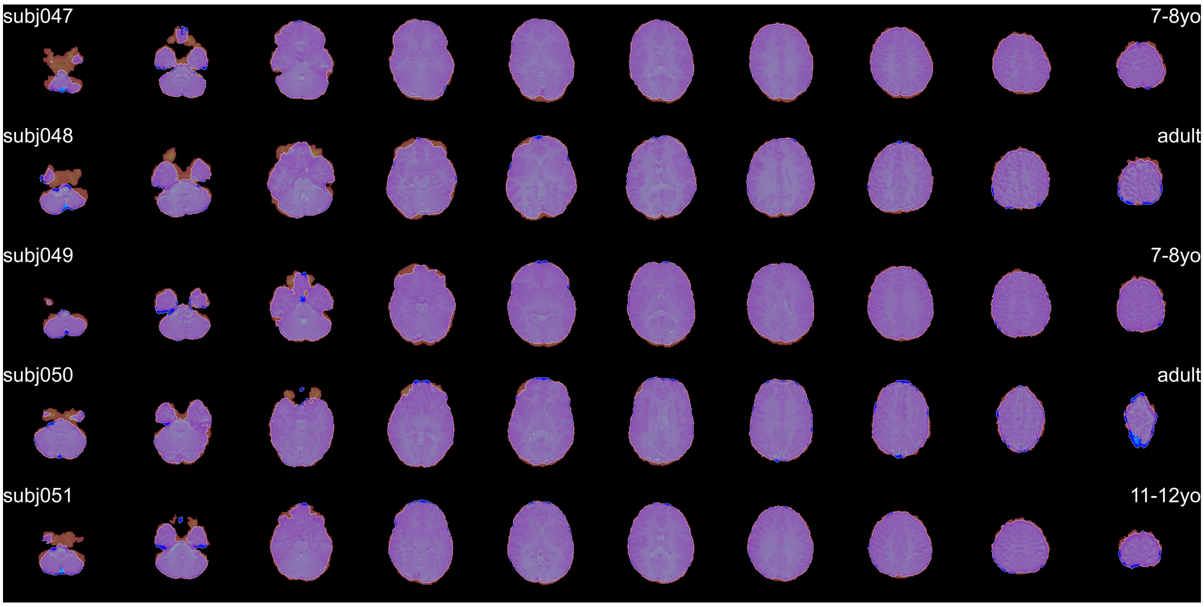


S2 Figure: Cross-modal alignment results of T1-weighed extracted brain image (red, transparent) onto diffusion-weighted extracted b0 brain image (blue with white outline). Ten (out of 60) slices are shown for each participant to demonstrate the alignment quality. Purple color indicates an overlap of T1 and DWI images. Solely blue or red voxels are mainly due to discrapancies in brain extractions, not due to misalignment, and therefore mostly appear in the most inferior or superior slices.
